# Reward modulates visual responses in the superficial superior colliculus of mice

**DOI:** 10.1101/2022.10.24.513513

**Authors:** Liad J. Baruchin, Matteo Alleman, Sylvia Schröder

**Affiliations:** School of Life Sciences, University of Sussex, Brighton BN1 9QG, UK; UCL Institute of Ophthalmology, University College London, London WC1E 6BT, UK; Center for Theoretical Neuroscience, Columbia University, NY, New York 10027, USA

## Abstract

The superficial layers of the superior colliculus (SC) are highly visual and receive direct input from the retina. Nonetheless, neural activity in the superficial SC (sSC) is modulated by locomotion and pupil-linked arousal. Here we show that visual responses of neurons in the sSC are additionally modulated by reward delivered prior to the visual stimulus. We trained mice to perform a visual detection task and recorded the activity of neurons in the SC using two-photon calcium imaging and electrophysiological recordings using high-density silicone probes (Neuropixels). Neurons across all layers of the SC responded to various task events, including reward delivery. However, responses to events like licking or movements did not explain the visual response modulation by reward. Electrophysiological recordings showed that most of the reward modulation occurred in the superficial rather than the deeper layers of the SC. Neurons also exhibited modulation by pupil-linked arousal, which was independent of the reward modulation. Performance of a population decoder to detect visual stimuli improved significantly by reward modulation but not by pupil-linked arousal modulation. Our results indicate that behavioural factors other than locomotion and arousal modulate visual activity in the SC.

## Introduction

The superior colliculus (SC) is a sensory-motor area in the mammalian midbrain, which receives visual input to its superficial layers (sSC) directly from the retina (Cang et al., 2018). Due to the strong retinal input, the sSC is considered mostly visual while the deeper layers of the SC (dSC) are thought to integrate this visual information to control orienting, attention and decision making (Basso and May, 2017; Felsen and Mainen, 2012; Jun et al., 2021; Wang et al., 2022, 2020; Zénon and Krauzlis, 2012). Visual processing in the sSC is modulated by locomotion and pupil-linked arousal (Ito et al., 2017; Joshi et al., 2016; Savier et al., 2019; Schröder et al., 2020), similar to the modulation by ongoing behaviour in the primary visual cortex (V1) (Vinck et al., 2015) and other brain areas (Kaplan and Zimmer, 2020). Here we show that, when mice are engaged in a visual detection task, visual responses in sSC neurons are additionally modulated by reward experienced in the previous trial.

## Results

We trained four mice to perform a visual detection task (**Figure 1A-C**). In each trial, a grating stimulus of varying contrast appeared in the left or right visual field. After an auditory go cue, the mouse had to move the stimulus towards the centre of the visual field by turning a steering wheel, or to hold the wheel still if no stimulus was presented. After correct choices, the mouse was rewarded with a drop of water, while incorrect choices were followed by auditory white noise.

**Figure 1.**
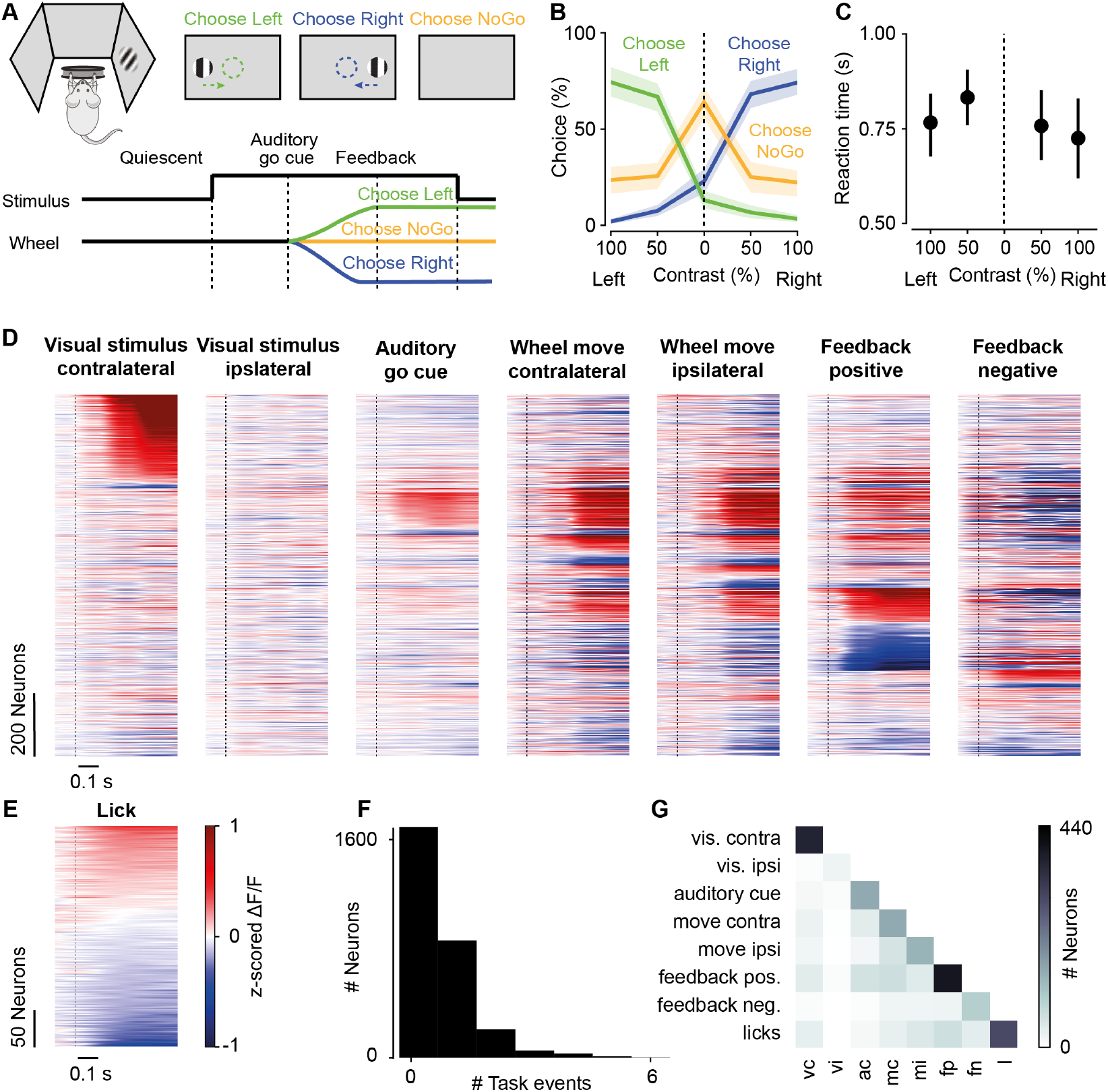
Superficial superior colliculus neurons respond to various task events. **A**. Top left: Experimental setup. Top right: Mouse has to choose left (right) when visual stimulus is presented on the left (right) and has to choose NoGo when no stimulus is presented. Bottom: Time course of each trial. Trial starts when wheel is kept still (quiescent). Visual stimulus appears. After a delay, auditory go cue signals that stimulus can be moved using wheel. Choice is recorded either when stimulus reaches decision threshold (at or away from target position), or at fixed delay after go cue. At time of choice, feedback is delivered: water reward for correct choices, auditory noise for incorrect choices. After a delay, visual stimulus disappears and new trial can be initiated. **B**. Psychometric curve. Percentage of left (green), right (blue), or NoGo (yellow) choices (mean ± SEM, 16 sessions, 4 mice) depending on stimulus contrast. Average performance, i.e., percentage of correct choices, across 16 sessions: 65.3 ± 0.3%; across 4 mice: 64.4 ± 0.2%. Trials in which the animal was disengaged, i.e., ≥3 consecutive NoGo trials, were discarded. **C**. Reaction time (mean ± SEM, 16 sessions) measured as time from go cue to time when stimulus reached its target. Only Go trials were considered. **D**. Mean calcium traces of 1,163 task responsive neurons (significant response to at least one task event; 2,855 neurons recorded) locked to onsets (dashed lines) of the visual stimulus (at 100% contrast), auditory go cue, wheel move, and feedback. Trace of each neuron was z-scored across session before averaging across trials. Order of neurons is the same across all plots. Neurons were first grouped by the first event they responded to (order of tested events as shown in plot), and then sorted by response amplitude. **E**. Mean calcium traces of 296 neurons (from 13 sessions) locked to onset of a lick outside the reward period. Only neurons with significant responses to licks shown. **F**. Distribution of number of task events eliciting significant responses. **G**. Number of neurons with significant responses to pairs of task events. Diagonal shows number of neurons responding to each task event.

Using two-photon imaging, we found that neurons in the sSC responded to several non-visual task events such as wheel movement or licking **(Figure 1D-G**). We recorded 2,855 neurons across 16 sessions from all four mice. 41% (1,163) of all neurons responded to at least one task event (termed *task responsive neurons*), while 11% (303) responded to more than one task event (**Figure 1F**). Task responsive sSC neurons exhibited significant responses to contralateral visual stimuli (367, 31%), ipsilateral stimuli (21, 2%), the auditory go cue (153, 13%), wheel movements in either direction (292, 25%), positive feedback/reward (394, 34%), negative feedback (96, 8%), and licking (296, 25%) (**Figure 1D,E**). Only few neurons that were responsive to reward also responded to wheel movements or licking (**Figure 1G**), indicating that bodily movements do not explain reward responses. Similarly, largely non-overlapping populations of sSC neurons responded to either wheel movements or licking (**Figure 1G**), indicating that sSC neurons differentiate between different actions. While these results confirm that sSC neurons are not purely visual, determining which events cause a response is difficult as they often occur near-simultaneously. We focused further analyses on responses to visual stimuli, which occur in relative isolation from other task events.

Visual responses of sSC neurons were most strongly affected by previous feedback and pupil-linked arousal (**Figure 2A-N**). We quantified tuning to visual contrast by fitting a hyperbolic ratio function (see *Contrast response fitting* in *Materials and Methods)* to stimulus responses for all neurons with significant responses to contralateral stimuli (**Figure 2A,B**). If this fit explained the data better than a flat line, the neuron was considered *visually responsive*. For these neurons, we determined the modulation of visual responses by four task variables: pupil-linked arousal (small or large pupil), previous feedback (positive or negative), upcoming action (Go or NoGo), and upcoming outcome (correct or incorrect). Examples of neurons modulated by arousal and previous feedback are shown in **Figure 2C-F**. Of all visually responsive neurons, 64% (204/319) were modulated by at least one task variable. Pupil-linked arousal affected 19% of neurons causing increases and decreases of visual responses (**Figure 2G,K**, **S1A**). Previous feedback affected a similar percentage of neurons (20%) with reward mostly increasing visual responses (**Figure 2H,L**, **S1B**). Only 14% (9/62) of these neurons with reward modulated visual responses also responded directly to positive reward, and none responded to negative reward. Finally, upcoming action (**Figure 2I,M**, **S1C**) or outcome (**Figure 2J,N**,**S1D)** modulated only very few neurons.

**Figure 2.**
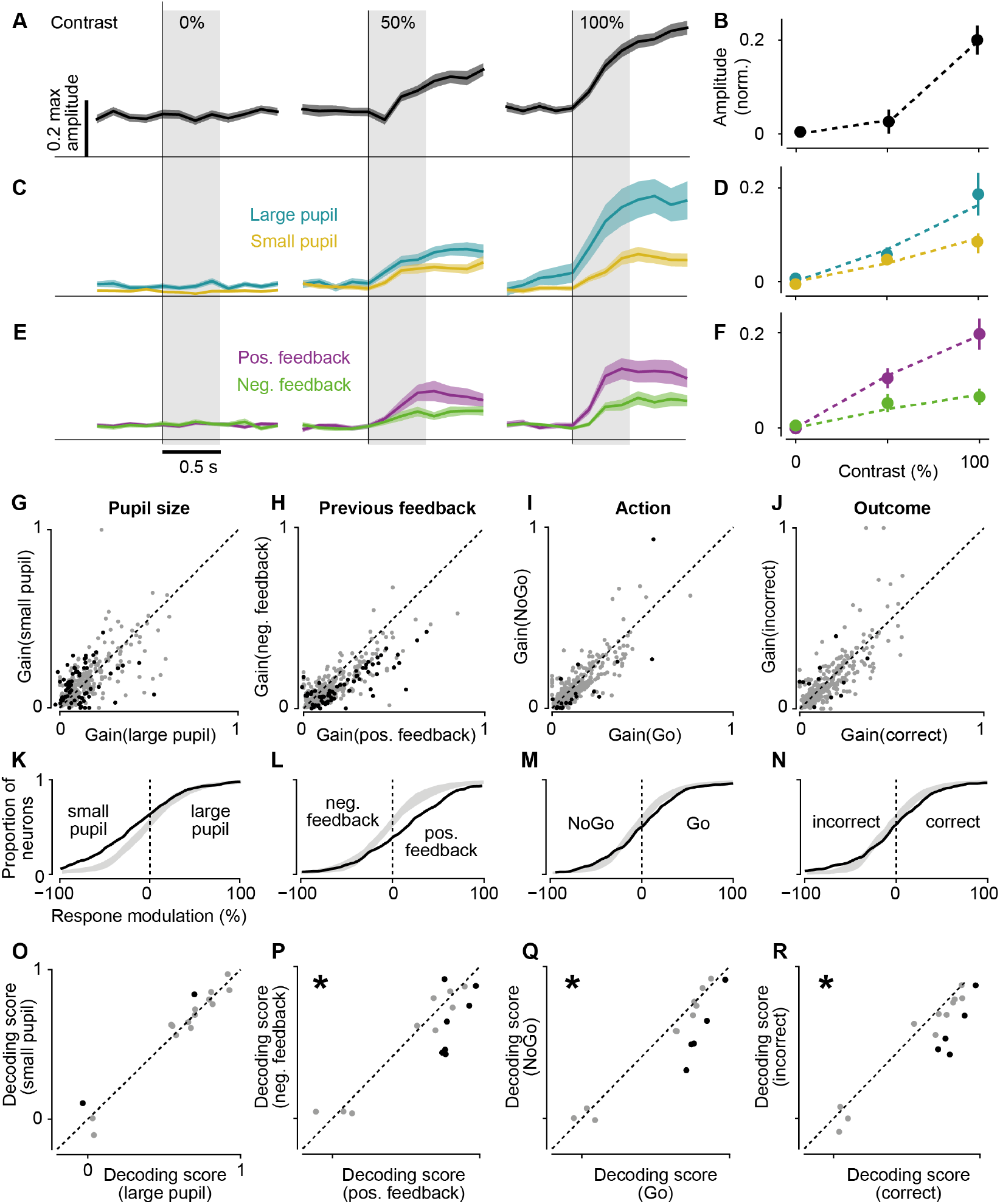
Visual responses in the superficial superior colliculus are modulated by task variables. **A**. Calcium traces (mean ± SEM) of one sSC neuron in response to visual stimuli of different contrasts. Vertical lines: stimulus onset; horizontal line at 0 ΔF/F. Shaded region: 0.0–0.5 s after stimulus onset, window used to determine response amplitudes (in **B**,**D**,**F**). **B**. Visual responses (mean ± SEM, same neuron as in **A**) fitted (dashed line) using a hyperbolic ratio function. Responses were baseline subtracted (mean of 0–0.5 s before stimulus onset). **C-F**. Same as in **A**,**B** for two different neurons. Trials were split according to pupil size (**C**,**D**) or previous feedback (**E**,**F**). **G-J**. Gain (R) of visually responsive neurons (319, 15 sessions, 4 mice) for pupil size (**G**), previous feedback (**H**), action (**I**), and outcome (**J**). Black dots: significantly different gains, p < 0.05, permutation test. Neurons with significantly increased/decreased responses: 21/41 during large pupil (**G**), 53/12 following reward (**H**), 14/10 during Go trials (**I**), 5/5 before correct choices (**J**). **K-N**. Cumulative distributions of response modulation (black line) by task variables. Grey shade: 2.5th to 97.5th interval of null distribution. **O-R**. Decoding scores of logistic regression models to detect presence of visual stimulus, tested on trials with different values of each task variable, e.g., small versus large pupil (**O**). Significant difference in predictive power per dataset (black dots) and across datasets (star) determined with permutation test (p < 0.05).

After rewarded trials, decoding the presence of a visual stimulus improved in populations of sSC neurons (**Figure 2O-R**). We trained a logistic regression decoder on the neuronal population of each session to detect contralateral stimuli based on peri-stimulus activity. The decoder was trained on a subset of trials and then tested on the held-out trials grouped by either pupil size, previous feedback, upcoming outcome, or action. Decoding performance was not influenced by pupil size (**Figure 2O**), but was improved by previous reward, correct choices, and Go actions (**Figure 2P-R**). The distributions of stimulus contrasts were similar across values of each task variable (**Figure S2A**). To account for the small differences, we balanced the data when training and testing the decoder so that stimulus contrast would not cause differences in decoding performance. Another confound of this analysis is that task variables may be correlated with each other. Animals were more likely to make a correct choice after a rewarded compared to an unrewarded trial (P(correct | positive feedback) = 0.665 ± 0.011, P(correct | negative feedback) = 0.588 ± 0.024, mean ± SEM, p < 0.01, t-test). Go actions were as likely after rewarded and unrewarded trials (P(Go | positive feedback) = 0.661 ± 0.041, P(Go | negative feedback) = 0.637 ± 0.031, p = 0.65). To control for the lack of independence between previous feedback, action, and outcome, we repeated the decoding analysis and determined the effect of each task variable while fixing another task variable to one value (**Figure S2B**). We found that decoding following reward was consistently better than following negative feedback using this control. In contrast, outcome and action had no effect on decoding performance when only trials following negative feedback were considered. To test whether previous reward had behavioural consequences, we used a logistic regression decoder to predict the outcome of a trial (correct or incorrect) based on stimulus contrast and outcome of the previous three trials. The results show that reward on the previous trial increased the likelihood of correct choice beyond the effect of stimulus contrast (**Figure S2C,D**).

Pupil-linked arousal and previous feedback were independent modulators of visual responses (**Figure 3A-C**). To test whether effects by arousal and reward had a common underlying cause, we compared pupil size during stimulus presentation after positive and negative feedback and found no difference (**Figure 3A,B**). In support of this conclusion, significant modulations by pupil size and previous feedback were independent of each other (**Figure 3C**).

**Figure 3.**
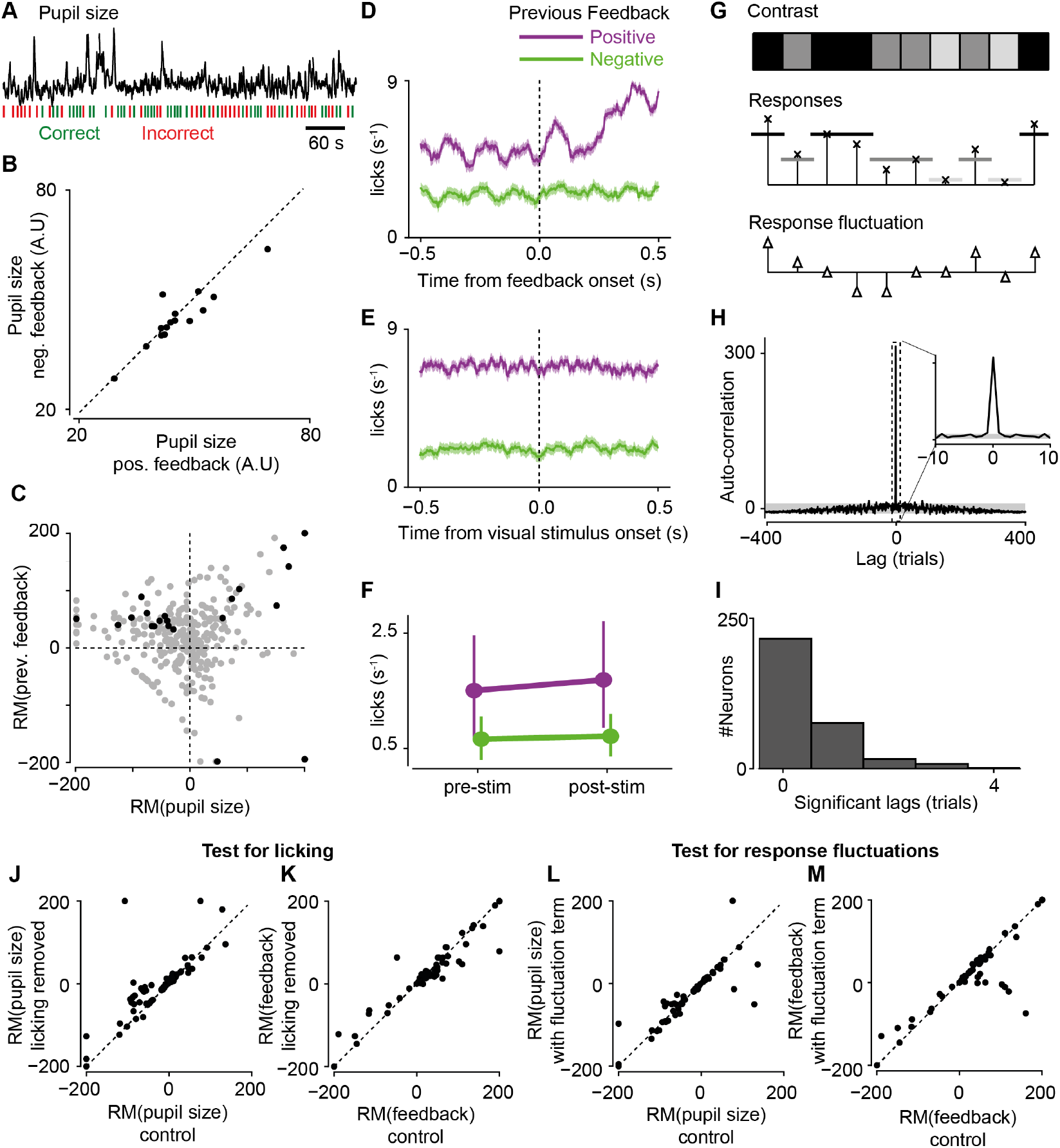
Modulation by previous feedback is independent of pupil size, licking, and response fluctuations. **A**. Example trace of pupil size and trial outcomes. **B**. Pupil size (mean across trials) following positive versus negative feedback (t(15) = 0.428, p = 0.675, paired t-test). **C**. Response modulation (RM) by pupil size versus previous feedback. Black dots: both RMs significant. Tests: all data: Pearson’s r = 0.13, p < 0.05, 319 neurons; black dots only: Pearson’s r = −0.07, p = 0.75, 21 neurons. **D**,**E**. Lick rate (mean ± SEM) locked to feedback onset (**D**) and stimulus onset following positive and negative feedback (**E**) for one session. **F**. Lick rate (mean ± SEM, 12 sessions) before (−0.5–0 s) and after (0–0.5 s) visual stimulus onset following positive or negative feedback. Mice licked more following rewarded than non-rewarded trials (F(1,44) = 9,543, p < 0.01), but lick rate was not significantly different between pre-and post-stimulus periods (F(1,44) = 0.0.23, p = 0.880). Test: two-way ANOVA (factors: time (pre-/post-stimulus) x feedback). **G**. Quantification of visual response fluctuation (see *Response fluctuation analysis* in *<u>Materials and Methods</u>*). **H**. Auto- correlogram of fluctuation trace (mean across visually responsive neurons in one session). Grey shade: 2.5th to 97.5th percentile interval of null distribution. **I**. Histogram of largest absolute lags with significant correlation strengths. **J**,**K**. Response modulation (RM) by pupil size (**J**) and previous feedback (**K**) for all trials (“control”) versus for no-lick trials (no licks in response window). RMs were not significantly different (pupil size: z = −0.05, p = 0.96, 62 neurons; feedback: z = 0.47, p = 0.64, 65 neurons; linear mixed-effects model). **L**,**M**. RM by pupil size (**L**) and previous feedback (**M**) as determined previously (“control”) and when accounting for response fluctuations. RMs were not significantly different (pupil size: z = 0.031, p = 0.98, 62 neurons; feedback: z = 0.44, p=0.66, 65 neurons; linear mixed effects model). **J-M**. Only neurons with significant RMs in control condition were considered (see **Figure 2G,H**).

Modulation by reward and pupil-linked arousal were independent of licking (**Figure 3D-F,J,K**). We found that reward induced a higher lick rate compared to negative feedback at the time of delivery, and that the lick rate after reward stayed high beyond the beginning of the next trial (**Figure 3D,E**). However, lick rate stayed constant before and after the onset of the visual stimulus (**Figure 3F**). Thus, licking was unlikely to induce larger visual responses, which were baseline subtracted. Similarly, lick rates were higher during large pupil trials but did not change around the time of stimulus onset (**Figure S3A**). To control for any effects related to licking, we repeated analyses while removing trials in which the animal licked during the tested time window (**Figure 3J,K**). This did not change the results regarding the modulation by reward.

Lastly, reward modulation was independent of response drift and specific to visual responses immediately following the reward (**Figure 3G-I,L,M**). Task engagement and motivation can affect both task performance and neural responses across the duration of an experimental session (Jacobs et al., 2020; Ortiz et al., 2020), which may explain the observed reward modulation. To quantify slow response fluctuations, we calculated the autocorrelation of stimulus-independent activity (**Figure 3G**). Hardly any neurons showed consistent response fluctuations exceeding two consecutive trials (**Figure 3H,I**). As a further test, we included stimulus-independent activity from the previous trial as additional predictor variable when fitting the contrast tuning curve. This did not significantly change the results (**Figure 3L,M)**. Finally, to test the time scale of reward effects, we modelled visual responses using positive and negative feedback delivered two trials prior to the visual stimulus instead of one trial prior. We found that the number of neurons modulated by feedback two trials before was largely reduced compared to feedback in the directly preceding trial (**Figure S3B**). This finding highlights the temporal specificity of reward modulation.

Reward and pupil modulation mostly affected neurons in the superficial rather than deep layers of the SC (**Figure 4**). Using Neuropixels probes, we recorded single neurons across the depth of SC in mice performing the a visual decision task similar to the one previously presented (see *Materials and Methods*) (6 mice, 15 sessions). The response patterns in the sSC (416 neurons recorded) and dSC (621 neurons recorded) were similar to those obtained using two-photon imaging (**Figure 4A,B, S4A-D)**. As before, visual responses tended to be positively modulated by previous reward, while pupil size led to increased and decreased responses (**Figure 4C,D**, **S4E,F**). Similarly to the two-photon recordings, only 6 neurons (17%) of the reward modulated neurons responded immediately to positive feedback, and none to negative feedback. While the distribution of visually responsive neurons was balanced across depth, the sSC contained 69% of all reward modulated neurons and 53% of all pupil modulated neurons (**Figure 4E**).

**Figure 4.**
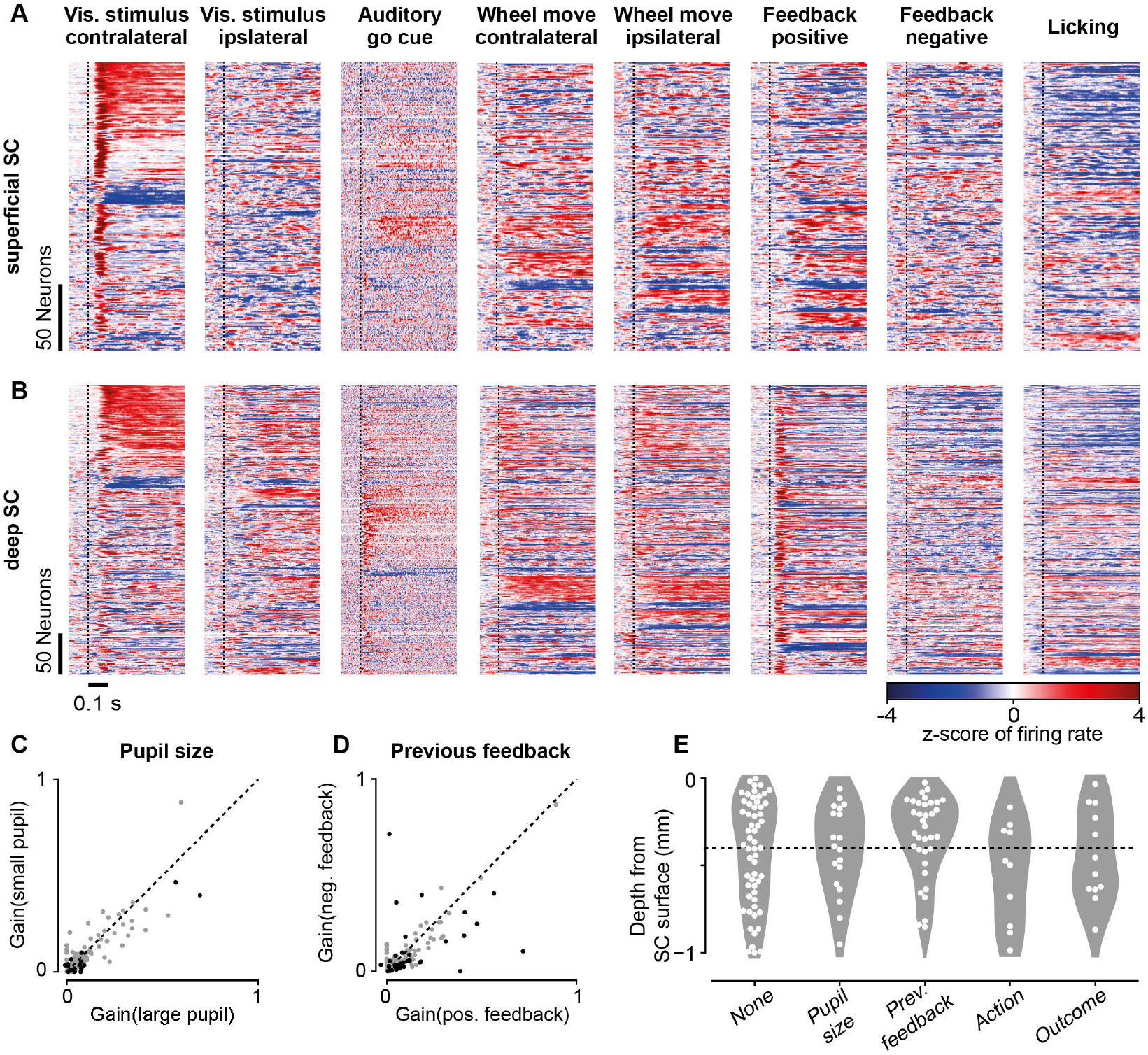
Responses to task events and modulation of visual responses by task variables extends to deep SC. **A**. Mean firing rates of 225 neurons (significant response to at least one task event; 416 neurons recorded, 15 sessions) in the sSC locked to onsets (dashed lines) of task events (for visual stimuli: only trials with 100% contrast on one side and 0% contrast on other side). Neurons significantly responded to contralateral stimuli (113, 50%), ipsilateral stimuli (21, 9%), auditory go cue (41, 18%), wheel movement (124, 55%), reward (82, 36%), negative feedback (13, 6%), and licking (79, 35%). **B**. As in **A**, but for 381 task responsive neurons (621 neurons recorded, 15 sessions) in the deep SC. Neurons significantly responded to contralateral stimuli (137, 36%), ipsilateral stimuli (63, 7%), auditory go cue (134, 35%), wheel movement (254, 67%), reward (178, 47%), negative feedback (49, 13%), and licking (155, 41%). **C**,**D**. Gain (R) of visually responsive neurons (158 neurons, 15 sessions, 6 mice) for pupil size (**C**), and previous feedback (**D**). Black dots: significantly different gains, p < 0.05, permutation test. Neurons with significantly increased/decreased responses: 13/5 during large pupil (**C**), 26/9 following reward (**D**). **E**. Depth of neurons within SC modulated by different task behaviours. Number of significantly modulated sSC/dSC neurons: 10/8 by pupil size, 24/11 by previous feedback, 4/10 by action, and 5/12 by outcome. Dashed line: border between superficial and deep superior colliculus at −400 μm.

## Discussion

We found that neurons in the SC responded to various task events and that reward modulated subsequent responses to visual stimuli. This modulation was independent of the modulation by pupil-linked arousal and appeared mostly in the superficial layers of the superior colliculus.

The SC has previously been shown to represent reward information (Felsen and Mainen, 2012; Ikeda and Hikosaka, 2003; Weldon et al., 2007), but modulation by reward in those studies was anticipatory as a result of learning, i.e. visual responses were affected before the expected reward was delivered, and was concentrated in the intermediated and deep layers of SC. Here instead we find that visual responses were modulated following reward delivery and that this modulation was exhibited by neurons across all layers of the SC and particular in the superficial layers.

We do not yet know the behavioural purpose of the modulation by reward. One possibility is that it serves a role in learning. Anticipatory reward modulation where stimuli associated with reward lead to larger responses has the advantage of increasing the neural decoding performance of behaviourally relevant sensory input. Such modulation has also been observed in other visual areas like V1 (Goltstein et al., 2018; Henschke et al., 2020; Jurjut et al., 2017). The retroactive reward modulation we observed here cannot fulfil the same purpose as it affects the response to a stimulus that in our task paradigm is irrelevant to the previously experienced reward. In a natural environment, however, sensory stimuli following reward may be highly relevant to associate the external circumstances with the reward. Thus, retroactive reward modulation can be beneficial depending on stimulus and reward statistics. A testable hypothesis then is that any reward, regardless of task, should increase visual responses in the SC.

The mechanism underlying the observed reward modulation is unclear. Potential candidates are other cortical areas, e.g., the cortical eye fields as shown in monkeys (Coe et al., 2002; Ikeda and Hikosaka, 2003; Leon and Shadlen, 1999). Another possibility is a contribution from serotonin, which encodes reward (Li et al., 2016) and specifically targets the sSC (Villar et al., 1988).

## Acknowledgments

We thank Profs Matteo Carandini (MC) and Kenneth D. Harris (KDH) for supervision and support of this project in their laboratory during data acquisition; we thank Miles Wells, Laura Funnell, Hamish Forrest, and Rakesh Raghupathy for help with training of animals and histology; we thank Charu Reddy for help with mouse husbandry; we thank Chris Burgess for help with setting up the behavioural paradigm and fixing various problems regarding experimental rigs and mouse training; we thank Michael Krumin for help with performing two-photon imaging experiments; we thank Nicholas A. Steinmetz for help with performing electrophysiological experiments; we thank Prof Miguel Maravall and Dr Benjamin Evans for feedback on the manuscript.

This work was supported by the People Programme (Marie Curie Actions) of the European Union’s Seventh Framework Programme (FP7/2007-2013) under REA Grant Agreement No 62387 (to SS), by the Biotechnology and Biological Sciences Research Council (grant BB/P003273/1 to MC and SS), by Wellcome Trust grants (095669 and 205093 to MC and KDH), and by a Sir Henry Dale Fellowship jointly funded by the Wellcome Trust and the Royal Society (grant 220169/Z/20/Z to SS). For the purpose of open access, the author has applied a CC BY public copyright licence to any Author Accepted Manuscript version arising from this submission.

## Author Contributions

Conceptualization: S.S., L.B.; animal training: M.A., S.S.; data collection: S.S., M.A.; data curation: S.S., M.A.; data analysis: L.B.; supervision: S.S.; writing: L.B., S.S.

## Data and code availability

The pre-processed data generated in this study are available at https://doi.org/10.25377/sussex.c.6208999; code used to analyse pre-processed data is available at https://github.com/liadJB/Baruchin-et-al-2022. The raw data are available on reasonable request.

## Materials and Methods

All procedures were conducted in accordance with the UK Animals Scientific Procedures Act (1986) under personal and project licenses released by the Home Office following appropriate ethics review.

### Animals

For two-photon imaging, we used 4 mice (3 male, 1 female) obtained by crossing Gad2-IRES-Cre (www.jax.org/strain/010802) and Ai9 (www.jax.org/strain/007909). The heterozygous offspring expressed TdTomato in glutamate decarboxylase 2-positive (GAD2+) cells. For electrophysiological recordings, we used 3 inbred C57Bl/6J mice (www.jax.org/strain/000664; 2 male, 1 female) and 3 mice (3 male) obtained by crossing PV-Cre (www.jax.org/strain/008069) and Ai32 (RCL-ChR2(H134R)/EYFP, www.jax.org/strain/012569). The heterozygous offspring expressed ChR2 in parvalbumin-positive cells, however, no optogenetic manipulations were performed.

Animals were 6-29 weeks old at the time of surgery with a mean weight of 27.4 g (20.0-43.2 g) and were used for experiments up to the age of 77 weeks. Mice were kept on a 12-h light: 12-h dark cycle. Most animals were single housed after the first surgery.

### Surgery

Animals were anesthetized with isoflurane (Merial) at 3.5% for induction, and 1-2% during surgery. Carprofen (5 mg/kg; Rimadyl, Pfizer) was administered subcutaneously for systemic analgesia, and dexamethasone (0.5 mg/kg; Colvasone, Norbrook) was administered as an anti-inflammatory agent to prevent brain swelling. The scalp was shaved and disinfected, and local analgesia (Lidocaine, 6 mg/kg, Hameln pharmaceuticals ltd) was injected subcutaneously under the scalp prior to the incision. Eyes were covered with eye-protective gel (Chloramphenicol, Martindale Pharmaceuticals Ltd). After the animal was placed into a stereotaxic apparatus (5% Lidocaine ointment, TEVA UK, was applied to ear bars), the skin covering and surrounding the area of interest was removed, and the skull was cleaned of connective tissue. A custom made headplate was positioned above the area of interest and attached to the bone with Superbond C&B (Sun Medical). Throughout all surgical procedures, the animal was kept on a heating pad to stabilize body temperature at 37°C. Subcutaneous injections of 0.01 ml/g/h of Sodium Lactate (Hartmann’s solution) were given. After the surgery, the animal was placed into a heated cage for recovery from anaesthesia. Mice were given three days to recover while being treated with Carprofen.

In animals used for two-photon imaging, a circular 4 mm craniotomy (centred at approximately 4.2 mm AP and 0.5 mm ML from Bregma) was made using a biopsy punch (Kai medical) and a fine-tipped diamond drill (Type 250-012 F, Heraeus). To reduce bleeding from the bone and from the dura we used bone wax and gel foam, and we cooled the area by applying cold cortex buffer. As the posterior SC is covered by a large sinus running through the dura, we permanently pushed the sinus anteriorly to gain optical access to the SC. We first made a cut into the dura directly posterior to the transverse sinus spanning the whole width of the craniotomy. Then we inserted a custom-made implant into the cut and pushed it anteriorly and a few 100 microns down to apply some pressure on the brain and thus stabilize the imaging. The implant was made of a short tube (2.29 mm inner diameter, 1 mm length) made of stainless steel (MicroGroup, Medway, Massachusetts). A 3 mm glass coverslip (#1 thickness, World Precision Instruments) was glued onto the tube to seal the end that was inserted into the craniotomy. A stainless-steel washer was glued onto the other end of the tube. The washer had an inner diameter that fit exactly around the tube and an outer diameter of 5 mm (Boker’s, Minneapolis, Minnesota). All three pieces were glued to each other using a UV curing adhesive (NOA61, Thorlabs). The glass coverslip was slightly larger than the outer diameter of the tube so that it could be slipped underneath the dura. The implant was placed into the craniotomy so that the washer was sitting on top of the skull and provided stability for the implant. The implant was fixed to the skull with Vetbond (3M) and Superbond C&B (Sun Medical). To prevent any dirt from staining the glass coverslip, we filled the tube of the implant with Kwik-Cast (World Precision Instruments), which could be easily removed before imaging.

For two-photon calcium imaging of activity in SC neurons, we injected the virus AAV2/1.Syn.GCaMP6f.WPRE.SV40 at a final concentration of 2.30-4.39e12 GC/ml after making the cut into the dura. 115-138 nl of the virus were injected 300-500 μm below the brain surface of the SC. The virus was injected at a rate of 2.3 nl every 6 s (Nanoject II, Drummond). The injection pipette was kept in place for about 10 min after the end of the injection.

For electrophysiological recordings, craniotomies were centred above the SC, −4.16 mm AP and 1.0 mm ML from Bregma, and performed the day before the first recording session (in one case 3 h before).

### Animal training

The training procedure has been described in detail previously (Burgess et al., 2017). After at least four days of recovery after surgery, mice were acclimatized to being handled and head-fixed on the experimental apparatus for at least three days. Before training, they were placed on a water control schedule, in which they received at least 40 ml/kg/day. During training, mice received about 3 μl of water at the end of each correct trial. After the daily training, they received top-up fluids to achieve the minimum daily rate. Body weight and potential signs of dehydration were monitored daily, and mice were given free access to water when they were dehydrated, or their weight fell below threshold. The training started with a simplified version of the task involving only high-contrast stimuli and longer response times. As performance improved, lower contrast stimuli were introduced and response times were reduced.

### Experimental apparatus

The experimental apparatus was described in detail before (Burgess et al., 2017). The mouse was head-fixed and surrounded by three computer screens (for two-photon imaging: Iiyama ProLite E1980SD placed ∼20 cm from the mouse’s eyes; for electrophysiology: Adafruit, LP097QX1 placed ∼11 cm from the mouse’s eyes; 60 Hz refresh rate for both models) at right angles covering approximately 270 × 70 degrees of visual angle. In some experiments, Fresnel lenses (BHPA220-2-6 or BHPA220-2-5, Wuxi Bohai Optics) were mounted in front of the monitors to compensate for reduction in luminance and contrast at steeper viewing angles relative to the monitors. In some of these experiments, lenses were coated with scattering window film (frostbite, The Window Film Company) to prevent specular reflections. A wheel was placed below the mouse’s forepaws and a rotary encoder (Kübler) measured the rotational movements of the wheel. A plastic tube for water delivery was placed in front of the mouse’s snout and calibrated amounts of water were released using a solenoid valve. Licking behavior was monitored by attaching a piezo film (TE Connectivity, CAT-PFS0004) to the plastic tube and recording its voltage. A detailed parts list of the apparatus can be found at http://www.ucl.ac.uk/cortexlab/tools/wheel. To track pupil size, we illuminated one of both eyes with an infrared LED (850 nm, Mightex SLS-0208-A or Mightex SLS-0208-B). Videos of the eye were captured at 30 Hz with a camera (DMK 23U618 or DMK 21BU04.H, The Imaging Source) equipped with a zoom lens (Thorlabs MVL7000) and a filter (long-pass, Thorlabs FEL0750; or band-pass, combining long-pass 092/52×0.75, The Imaging Source, and short-pass FES0900, Thorlabs).

### Behavioural tasks

The task in the two-photon imaging experiments closely followed the paradigm described earlier (Steinmetz et al., 2019). To start a trial, the mouse had to keep the wheel still for 0.8-1.4 s (randomly chosen from uniform distribution). At initiation of 60-90% of all trials, a visual stimulus was presented at 95-115 deg either to the left or the right of the vertical meridian. The stimulus was a Gabor patch with vertical orientation (90 deg), sigma of 9 deg, spatial frequency of 0.1 cycles per deg and contrast of 12.5-100% (the range of presented contrasts depended on each animal’s task performance). In the rest of the trials, no stimulus was presented. After 0.5-1.2 s (randomly chosen from uniform distribution), an auditory go cue (8 kHz for 0.2 s) signalled that the position of the stimulus (if present) was now coupled to the wheel movement. The time count between stimulus onset and go cue was reset if the mouse moved the wheel during this period. After the go cue, the mouse had to either move the presented stimulus towards the vertical meridian (to 50-60 deg in the two-photon paradigm, to 30-80 deg in the electrophysiology paradigm), or, if no stimulus was presented, to hold the wheel still (more precisely: not cross the threshold for moving a stimulus to the target position). The mouse had to perform its choice within a fixed interval (“response time”) of 1.5 s after the go cue. After a correct choice, the mouse was given a water reward of 2-2.3 μl as soon as the stimulus reached its target position or at the end of the response time. The visual stimulus (if present) disappeared 1-1.3 s after reward delivery. After an incorrect choice, i.e. moving the stimulus by 50-60 deg towards the periphery, not reaching the target position within the response time, or moving the wheel too far while no stimulus was presented, an auditory white noise stimulus was played for 1 s, after which the visual stimulus (if present) disappeared. The mouse could then start the next trial. Trials of different contrast conditions were randomly interleaved. However, the same condition was repeated when the mouse responded incorrectly.

The task in the electrophysiology experiments was very similar to the task just described. The only differences were the following: (1) The mouse had to keep the wheel still for 0.2-0.5 s to start a trial. (2) Two Gabor patches were presented simultaneously, one to the left and one to the right, at 26-80 deg from the vertical meridian. The mouse had to choose the patch with the higher contrast. (3) The width of the Gabor patches (sigma) varied between 8-11 deg and contrasts varied between 25-100%. (4) Any wheel movements between stimulus onset and go cue were simply ignored. (5) The go cue appeared 0.4–0.8 s after stimulus onset. (6) The negative feedback (auditory white noise) was played for 1.5-2 s. (7) The response time was 1.3-1.6 s long.

### 2p imaging

Two-photon imaging was performed using a standard resonant microscope (B-Scope, ThorLabs Inc.) equipped with a 16x, 0.8 NA water immersion objective (N16XLWD-PF, Nikon) and controlled by ScanImage 4.2 (Pologruto et al., 2003). Excitation light at 970-980 nm was delivered by a femtosecond laser (Chameleon Ultra II, Coherent). Multi-plane imaging was performed using a piezo focusing device (P-725.4CA PIFOC, Physik Instrumente, 400 μm range). Laser power was depth-adjusted and synchronized with piezo position using an electro-optical modulator (M350-80LA, Conoptics Inc.). The imaging objective and the piezo device were light-shielded using a custom-made metal cone, a tube, and black cloth to prevent contamination of the fluorescent signal caused by the monitors’ light. Emission light was collected using two separate channels, one for green fluorescence (525/50 nm emission filter) capturing the calcium transients and one for red fluorescence (607/70 nm emission filter) capturing the expression of TdTomato in inhibitory neurons of Gad-Cre x TdTomato mice.

For imaging neurons in SC, we used 2-4 imaging planes separated by 30-55 μm at depths of 12-260 μm from the surface of SC. The field of view spanned 410-770 μm in both directions at a resolution of 512 × 512 pixels. The frame rate per plane was 6-10 Hz. For one dataset, a single plane was captured at a frame rate of 30 Hz.

### Electrophysiology

Recordings were made using Neuropixels 1.0 electrode arrays (Jun et al., 2017). Probes were mounted to a custom 3D-printed piece and affixed to a steel rod held by a micromanipulator (uMP-4, Sensapex Inc.). Prior to insertion, probes were coated with a solution of DiI (ThermoFisher Vybrant V22888 or V22885) or DiO (ThermoFisher Vybrant V22886) by holding 2 μl in a droplet on the end of a micropipette and touching the droplet to the probe shank, letting it dry, and repeating until the droplet was gone. Probes had a soldered connection to short external reference to ground; the ground connection at the headstage was subsequently connected to an Ag/AgCl wire positioned on the skull. The craniotomies and the wire were covered with saline-based agar. The agar was covered with silicone oil to prevent drying. In some experiments a saline bath was used rather than agar. Probes were advanced through the agar and the dura, then lowered to their final position at ∼10 μm/sec. Electrodes were allowed to settle for ∼10 min before starting recording. Recordings were made in external reference mode with LFP gain = 250 and AP gain = 500. Data were filtered in hardware with a 300 Hz 1-pole high pass filter and digitized at 30 kHz. Recordings were repeated at different locations on each of multiple subsequent days.

### Pre-processing of imaging data

All raw two-photon imaging movies were analysed using suite2p (implemented in MATLAB, Mathworks) to align frames and detect regions of interest (Pachitariu et al., 2016b). We used the red channel representing TdTomato expressed in all inhibitory neurons to align frames, which yielded better results than alignment using calcium dependent fluorescence. Every aligned movie was inspected manually to check for failures in automatic alignment. Failures were corrected using different parameter settings where possible. Misaligned movie frames were discarded and ROIs in unstable regions of the field of view were not considered for further analysis.

Using the aligned movies and detected ROIs resulting from suite2p analysis, we extracted the fluorescence from the green and the red channel within each ROI. To correct the calcium traces for contamination by surrounding neuropil, we also extracted the fluorescence of the surrounding neuropil for each ROI using the green channel. The neuropil mask resembled a band surrounding the ROI with its inner edge having a distance of 3 microns away from the edge of ROI and its outer edge having a distance of 30 microns from the edge of the ROI. Pixels belonging to other ROIs were excluded. To correct for contamination, the resulting neuropil trace, N, was subtracted from the calcium trace, F, using a correction factor α: F_c_(t) = F(t) - α·N(t). The correction factor was determined for each ROI as follows. First, F and N were low-pass filtered using the 8th percentile in a moving window of 180 s, resulting in F_0_ an N_0_. The resulting traces F_f_(t) = F(t)-F_0_(t) and N_f_(t) = N(t)-N_0_(t) were then used to estimate α as described previously (Dipoppa et al., 2018). In short, N_f_ was linearly fitted to F_f_ using only time points when values of F_f_ were relatively low and thus unlikely to reflect neural spiking. F_c_ was then low-pass filtered as above (8th percentile in a moving window of 180 s) to determine F_c,0_. These traces corrected for neuropil contamination were then used to determine ΔF/F = (F_c_(t) - F_c,0_(t)) / max(1, meant(F_c,0_(t)).

To correct for potential brain movements, we used the red fluorescence traces of each ROI to regress out changes in fluorescence that were not due to neural activity. First, we low-pass filtered the red trace of each ROI (8th percentile in a moving window of 180 s) and subtracted it from the unfiltered trace to remove slow drifts and bleaching effects. Second, we applied a median filter to the resulting red trace (moving median in window of 10 s). Third, this trace was regressed out of ΔF/F.

We avoided sampling the same neurons more than once. We detected ROI pairs that were close to each other in neighbouring imaging planes and that had highly correlated calcium traces (ρ > 0.4, correlation between traces filtered using a moving median in a window of 5 samples). Only the ROI of each pair with the highest signal-to-noise ratio was used for further analyses. ROIs that had very long-lasting calcium transients (> 25 s) were removed.

### Spike sorting and firing rates

Extracellular voltage traces were pre-processed using common-average referencing: subtracting each channel’s median to remove baseline offsets, then subtracting the median across all channels at each time point to remove artifacts. Electrophysiological data collected in SC was spike sorted using Kilosort2 (available at https://www.github.com/MouseLand/Kilosort2) with standard parameters (Pachitariu et al., 2016a). After sorting, all automatically-detected spike clusters were curated manually using phy (https://github.com/kwikteam/phy). The spiking data of each neuron was then converted into a firing rate using a 5 ms window.

### Depth of electrophysiological recordings

We used the LFP recorded on each channel of the Neuropixels probes to determine the surface of the SC (Ito et al., 2017). We used LFP responses to the following visual noise stimulus: We presented white squares of 8 visual degrees edge length, positioned on a 10 × 36 grid (some stimulus positions were located partially off-screen) on a black background. The presentation of each white square lasted on average 6 monitor frames (100 ms), and their times of appearance were independently randomly selected to yield an average rate of approximately 0.12 Hz. We then determined the stimulus square that elicited the most negative LFP amplitudes based on the average LFP waveform triggered by a change of luminance in each square. At the time of peak negative LFP amplitude measured on the probe channel with the most negative peak, we calculated the LFP amplitude as a function of depth for all probe channels separately. We then determined the depth at half height of the negative peak (located below the depth of the peak) for all right and left channels on the probe. The average result of the right and left probe channels was defined as the surface of the SC. We defined the depth of 400 μm below the SC surface as the border between superficial and deep SC (Ito et al., 2017).

### Pupil Tracking

We used DeepLabCut (Mathis et al., 2018) to detect 8 landmarks in our eye videos: top/bottom/left/right edge of the pupil and top/bottom/left/right edge of the eye lids. The trained network is available from https://github.com/sylviaschroeder/PupilDetection_DLC. Pupil diameter was defined as the distance between the top and bottom edges (height) of the pupil. This measure was chosen as it is relatively independent from horizontal eye movements (the most common eye movements mice perform), while the distance between left and right pupil edges (width) changes with eye movements. For time points when the lids covered top or bottom pupil edges (i.e. certainty of top/bottom markers returned by DLC was low or the markers were very close to markers for top/bottom edges of the lid), we used eye position and left and right pupil edges to estimate pupil diameter. Based on samples where the pupil was not covered by the lid, we used local linear regression (Matlab function fit, with method lowess and span = 0.1) to fit pupil height given pupil width and horizontal position of pupil center. Times when the eye was closed, including 5 frames before and after detected closure, were ignored for further analysis. The eye was defined closed when one of the following criteria was true: (1) certainties of left or right pupil marker was low, (2) small distance between top and bottom lid, (3) certainties of top and bottom pupil marker was low, or (4) lid too close to centre of pupil. Values of pupil diameter were median filtered with a span of 5 frames. Pupil size per trial was determined by averaging pupil size in a −0.5–0.5 s window from stimulus onset. A trial was considered a large-pupil trial if the average value was above the median pupil size for the session.

### Responsiveness to task events

We tested responses to the following events: (1) contralateral visual stimulus in trials without wheel movements within test window (to dissociate visual from movement responses), (2) ipsilateral visual stimulus in trials without wheel movements, (3) auditory go cue in trials without wheel movements (to dissociate auditory from movement responses), (4) ipsilateral and contralateral wheel movements in no-stimulus trials (to dissociate wheel from visual movement responses), (5) positive and negative feedback in no-stimulus trials (to dissociate feedback from visual responses), and (6) licks at least 0.5 s outside the reward delivery time to dissociate licking from reward responses. In the case of lick bursts (licks within less than 0.3 s from each other), only the first lick was taken into consideration. We measured average calcium responses or firing rates in windows of 0–500 ms after event onset (post-event responses) and compared these to average responses in windows of 0–500 ms before event onset (pre-event responses) using paired t-tests. For responses to visual stimuli (contra-or ipsilateral, respectively), we used a two-way ANOVA with pre- and post-event time as one factor and stimulus contrast as second factor. A neuron was considered responsive to a task event if p < 0.05, after Bonferroni multiple test correction.

### Contrast response fitting

Responses to visual stimuli were determined by averaging neuronal activity across a window of 500 ms after stimulus onset for two-photon recordings and 200 ms after stimulus onset for electrophysiology recordings. The time window was chosen based on the peak latency of the visual response recorded by the two methods. These responses were used to fit a hyperbolic ratio function (Albrecht and Hamilton, 1982):

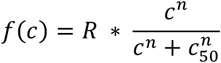

where R is gain, c is stimulus contrast, c_50_ is contrast at half-saturation, and n is rate of change. For the electrophysiological data where two Gabor patches were presented simultaneously only the contrast at the contralateral side was taken into account while the ipsilateral contrast was ignored because ipsilateral stimuli did not influence visual responses. The curve was only fit to neurons that exhibited a significant difference in response amplitude after stimulus onset compared to baseline activity before stimulus onset (see *Responsiveness to task events*). If the fit resulted in an explained variance larger than that of flat line fitted to the responses, the neuron was considered *visually responsive*. 48 neurons in the two-photon recordings and 92 neurons in the Neuropixels recordings had significant responses to contralateral visual stimuli but could not be fit well with the hyperbolic ratio function. Only visually responsive neurons were tested for modulation by task variables. To test for the effect of pupil-linked arousal (small or large pupil), previous feedback (reward or negative feedback), upcoming action (Go or NoGo), and upcoming outcome (correct or incorrect) on response amplitude, we fitted the individual neuronal responses to the same function as above, but modified the gain as follows:

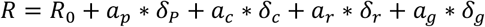

Where the factors represented by *a* are weights and δ ∈ {0, 1} represented the presence or absence of a specific task variable, e.g., correct outcome, in each trial. To find the best model explaining the visual responses, we fitted variations of this function, leaving out any possible combination of δ*s* so that either all or only some of the behavioural conditions would be considered for the modulation of visual responses. The function with the highest cross-validated explained variance determined which task variables had significant impact.

The cross-validated explained variance was calculated using a 10-fold cross-validation, whereby the dataset was divided into 10 parts, and the model trained on 9 of those, while the remaining dataset was used to test the fit. To verify the choice of model, we employed a permutation test (see *Permutation Test*) and only used models that resulted in p-values smaller than 0.05.

### Response modulation

Response modulation of visually responsive neurons was calculated using the following formula:

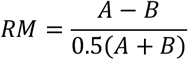

where *A* was the gain in either large pupil, previously rewarded, Go, or correct trials; and *B* was the gain in either small pupil, previously negative feedback, NoGo, or incorrect trials.

### Control for licking

We controlled for licking by removing trials where the mouse licked during the response window used to measure the visual response (0–500 ms after stimulus onset). The contrast response fitting was then performed on the rest of the trials as described above.

### Response fluctuation analysis

Response fluctuations of neural response can occur across long timescales of many seconds to minutes. To measure response fluctuations across a recording session, we accounted for response variation due to the different visual contrasts in the following way. For each neuron, response amplitudes to each visual stimulus were calculated. The time windows were the same as those described in *Contrast response fitting*. For each contrast, single trial responses were z-scored by the contrast specific mean and standard deviation. To measure the time scale of fluctuations, we calculated the auto-correlogram of this fluctuation signal. To test if any point in the auto-correlogram is significantly larger than expected, we generated a null distribution by randomly drawing 500 values from the auto-correlogram. We used the 2.5th to 97.5th percentile interval of the null distribution to determine significant values as consecutive lags (from 0) that were outside the interval.

### Control for response fluctuations

To control for response fluctuations, i.e., trial-to-trial variability, the tested function was modified to include a signal fluctuation variable, as follows:

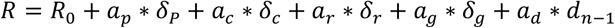

where *d* is the fluctuation signal of the previous trial, calculated as explained in *Response fluctuation analysis*.

### Decoding stimulus presence

We used responses, averaged across a 0.5 s time window after stimulus onset, of all neurons in the recorded population to decode the presence or absence of a contralateral visual stimulus. We trained a logistic regression model on 50% of the data, and used the other 50% to test the model (Christensen and Pillow, 2022). The fitted function was:

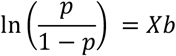

where p is the probability that stimulus is present, X is the neuronal response matrix [trial x number of neurons], and b is the coefficient vector [number of neurons x 1].

We split the test data into two groups. The split was either according to large/small pupil, previously positive/negative feedback, Go/NoGo, or correct/incorrect. We stratified the data so that both splits were of equal size. We repeated the procedure 10 times and used the mean prediction score of each group for comparison. We scored the prediction using the Matthew Correlation Coefficient (Matthews, 1975). The score ranges from −1 to 1, where 0 is chance level, 1 is a perfect prediction, and −1 is a perfectly opposite prediction. This measure combines prediction accuracy and recall. To control for possible covariance between the task variables (pupil size, previous feedback, action, and outcome), we repeated the procedure described above while keeping one variable constant (e.g., by only considering trials where pupil size is large).

### Predicting trial outcome

To predict trial outcome, we used a logistic regression model with 7 independent variables: an intercept, a stimulus of 0% contrast on both sides (*Zero* ∈ {0, 1}), a stimulus with left contrast > 0% (*Left* ∈ {0, 0.125, 0.25, 0.5, 0.75, 1}), a stimulus with right contrast > 0% (*Right* ∈ {0, 0.125, 0.25, 0.5, 0.75, 1}), and correct choice on each of the previous 3 trials (*R*_*T-1*_, *R*_*T-2*_, *R*_*T-3*_ ∈ {0, 1}). The model was trained on 80% of the data and tested on the remaining 20%. The formula was:

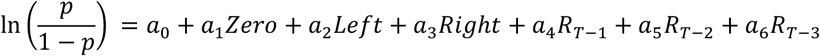

where a_0_-a_6_ are the coefficients and the variables are as described above.

Like for the logistic regression to predict stimulus presence (see previous paragraph), we used the Matthew Correlation Coefficient (Matthews, 1975) to score the model prediction. To measure the significance of the parameter coefficients, we performed a Wald test (Wald, 1943) with the null hypothesis of the parameter being equal to 0.

### Permutation Test

Permutation tests were performed by generating surrogate datasets from the recorded data, computing the test statistic for each surrogate dataset, and comparing the resulting null distribution to the test statistic of the recorded data. Surrogate datasets were generated by randomly permuting the values of the task variables, e.g., small and large pupil size.

To determine significance of gain changes in the contrast response function due to task variables (**Figure 2G-J, Figure 3C,J-M, Figure 4C,D, Figure S2B**), we generated 200 surrogate datasets per recorded dataset by permuting the values of the task variable (e.g. small and large pupil). For each neuron, we fitted the hyperbolic ratio function to each of the permuted datasets to generate a null distribution of 200 instances of cross-validated (10-fold) variance explained. We used the same fits to the surrogate datasets to derive the null distribution of response modulations and to determine the 2.5^th^ to 97.5^th^ percentile interval of null distribution for each possible value of response modulation (**Figure 2K-N, Figure S3E,F**).

To determine significance for the performance of the logistic regression model that predicts stimulus presence (**Figure 2O-R, Figure S1C**), we generated 200 surrogate datasets per recorded dataset by permuting the values of the task variable (e.g. previous positive and negative feedback). The test statistic was the difference between prediction scores for the two splits of the test data (according to value of the task variable). In order to determine significance across datasets, the mean across sessions of the actual test statistic was then compared to the distribution of test statistics across the surrogate datasets.

To determine significance for the logistic regression model that predicts behavioural outcome (**Figure S2C,D**), we generated 500 surrogate datasets per recorded dataset by permuting the value of the trial outcome, i.e. correct and incorrect choices. Only scores larger than the 95^th^ percentile of the surrogate scores were considered significant.

## Supplementary Material

**Figure S1.**
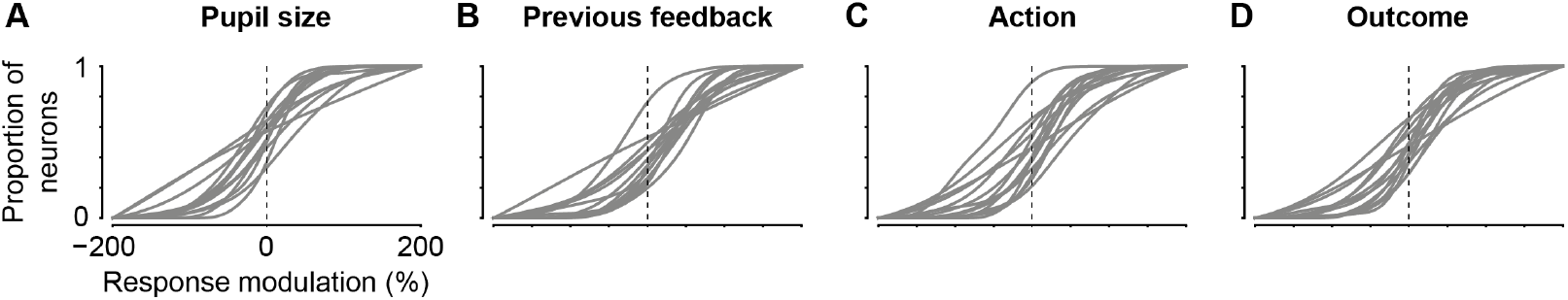
Modulation by task variables in single datasets. **A-D**. Cumulative distributions of response modulation by task variables for each of the 16 sessions. Note that one session has a flat distribution since only two visually responsive neurons were found within the session.

**Figure S2.**
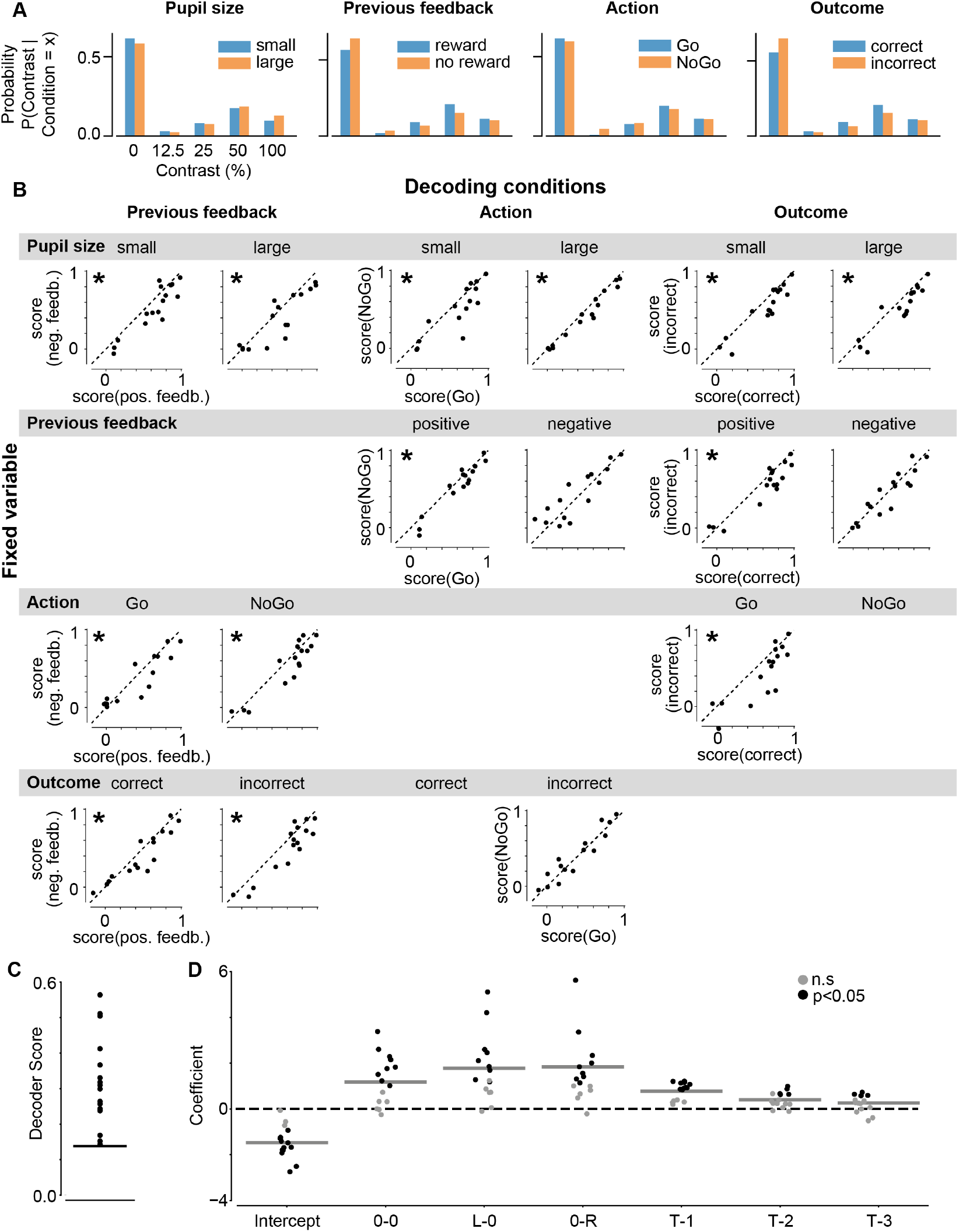
Control for interaction of task variables when decoding stimulus presence. **A**. Probability of contralateral stimulus contrasts presented across all trials for both values of each task variable, e.g., small and large pupil (first column). **B**. Scores of logistic regression models trained to detect the presence of the visual stimulus contralateral to the recorded SC neurons. Models were tested under both values of each task variable (columns), e.g., previous positive versus negative feedback (left column), while a second task variable was fixed (rows), e.g., small or large pupil (first row). Significant score differences across sessions marked by star (p < 0.05, permutation test). Note, no visual stimuli were presented in correct NoGo trials, therefore the decoder could not be fit in combinations of these conditions. **C**. Scores of logistic regression models trained to predict trial outcome based on visual stimulus in the current trial (0-0, L-0, or 0-R) and outcome of the previous three trials (T-1, T-2, T-3). Line indicates the score equivalent to p = 0.05 (permutation test). All trained models performed better than chance (n = 16 sessions). **D**. Regression coefficient values for each trained model. Black: coefficients are significantly different from 0 (p < 0.05, Wald test). Grey lines show medians of coefficients across sessions. In 10/16 sessions, reward in the previous trial increased the likelihood of correct outcome (see “T-1”).

**Figure S3.**
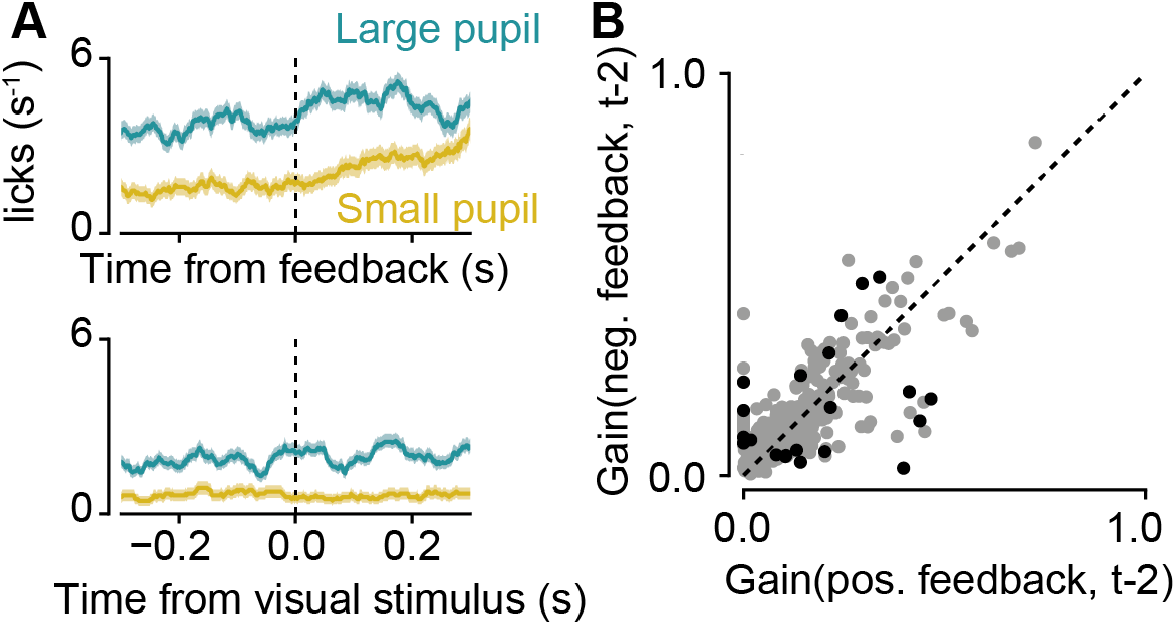
Control for licking and feedback history. **A**. Lick rate (mean ± SEM) locked to feedback delivery before small and large pupil trials (top) and locked to visual stimulus onset during large and small pupil trials (bottom) for one example session. Mice licked more during large pupil trials (F(1,44) = 4.02, p < 0.05), and after feedback delivery (F(1,42) = 6.55, p < 0.05). They did not lick more after stimulus presentation (F(1,44) = 0.131, p = 0.720). Test: two-way ANOVA (factors: time (pre-/post-event) x pupil size (large/small)). **B**. Gain (R) for positive versus negative feedback two trials before the current trial (281 neurons, 15 sessions, 4 mice; black dots: significantly different gains, p < 0.05, permutation test).

**Figure S4.**
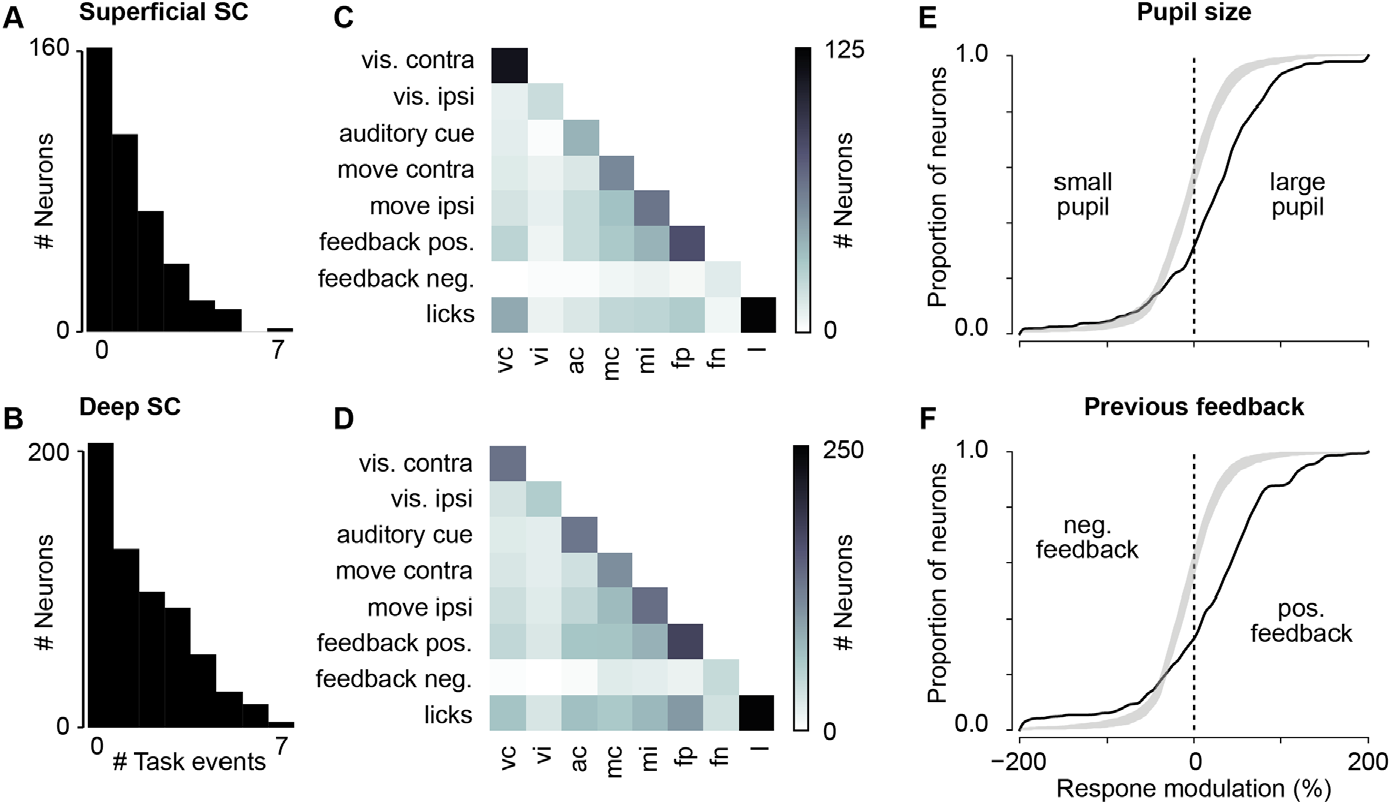
Task modulation across all layers of the SC. **A**,**B**. Distribution of number of task events eliciting significant responses in sSC neurons (**A**; 54% responded to at least one task event, and 30% responded to more than 1 task event), and dSC neurons (**B**; 61% responded to at least one task event, 42% responded to more than 1 task event). **C**,**D**. Number of sSC neurons (**C**) and dSC neurons (**D**) with significant responses to pairs of task events. Diagonal shows the number of neurons responding to a single task event. **E**,**F**. Cumulative distribution of response modulation (black line) by pupil size (**E**) and previous feedback (**F**) for all visually responsive neurons. 2.5th to 97.5th intervals of null distribution shown by grey shade.

